# Microbiome signatures of virulence in the oral-gut-brain axis influence Parkinson’s disease and cognitive decline pathophysiology

**DOI:** 10.1101/2024.10.14.618175

**Authors:** Frederick Clasen, Suleyman Yildirim, Muzaffer Arıkan, Fernando Garcia-Guevara, Lütfü Hanoğlu, Nesrin H. Yılmaz, Aysu Şen, Tuğçe Kahraman Demir, Zeynep Yıldız, Adil Mardinoglu, Mathias Uhlen, Saeed Shoaie

## Abstract

The human microbiome is increasingly recognized for its crucial role in the development and progression of neurodegenerative diseases. While the gut-brain axis has been extensively studied, the contribution of the oral microbiome and gut-oral tropism in neurodegeneration has been largely overlooked. Cognitive impairment (CI) is common in neurodegenerative diseases and develops on a spectrum. In Parkinson’s Disease (PD) patients, CI is one of the most common non-motor symptoms but its mechanistic development across the spectrum remains unclear, complicating early diagnosis of at-risk individuals. Here, we generated 228 shotgun metagenomics samples of the gut and oral microbiomes across PD patients with either mild cognitive impairment (PD-MCI) or dementia (PDD), and a healthy cohort, to study the role of the gut and oral microbiomes on CI in PD. In addition to revealing compositional and functional signatures, the role of pathobionts, and dysregulated metabolic pathways of the oral and gut microbiome in PD-MCI and PDD, we also revealed the importance of oral-gut translocation in increasing abundance of virulence factors in PD and CI. The oral-gut virulence was further integrated with saliva metaproteomics and demonstrated their potential role in dysfunction of host immunity and brain endothelial cells. Our findings highlight the significance of the oral-gut-brain axis and underscore its potential for discovering novel biomarkers for PD and CI.

## INTRODUCTION

Neurological disorders are the leading cause of physical and cognitive disability around the world, currently affecting approximately 15% of the worldwide population and expected to increase in future decades due to an ageing population, industrialization and changes in environmental impacts ^1–3^. Parkinson’s Disease (PD) is a complex neurodegenerative disease with the fastest growing prevalence worldwide ^2,4^. While it is primarily characterized by motor symptoms such as involuntary shaking, slow movements, and muscle stiffness, one of its most common non-motor dysfunctions is cognitive impairment (CI). There is a high risk of dementia in patients with PD with nearly half of patients reaching the dementia stage within 10 years after diagnosis and virtually all patients develop full dementia within 20 years after diagnosis ^5^. CI develops on a spectrum that ranges from mild cognitive impairment (MCI) to full-scale dementia ^4,6,7^. Identification of the risk of developing CI and cognitive decline are important for clinical management of NDs ^5^. However, the evaluation of cognition remains challenging and there is currently an unmet need on whether patients with neurodegenerative diseases have CI or are at risk for further cognitive decline. Non-genetic factors, such as microbiome and environmental impacts, including diet, pollution, and drugs exposure, may have a significant role in this ^8,9^.

A growing body of evidence links the gastrointestinal (GI) tract with neurodegenerative diseases, including PD, and GI dysfunction is common in patients with PD ^10,11^. As such, several studies have investigated the role of the gut microbiome in PD for novel diagnostic and treatment avenues as well as a better understanding of the gut-brain axis ^12^. Several studies consistently indicated an increased abundance in *Akkermansia, Bfidobacterium* and *Lactobacillus*, and a depletion in butyrate producers such as *Roseburia, Faecalibacterium* and *Blautia* in PD patients ^2^. One of the key mediating factors of the gut microbiome composition is microbial metabolites and virulence that can have an impact on PD and CI onset and progression. This could be through induction of neuroinflammation and oxidative stress that exacerbate neurodegeneration ^13,14^. Among microbial metabolites, short chain fatty acids (SCFAs) production and especially butyrate has shown to have neuroprotective effects ^15^. At the same time the secretion of bacterial endotoxins and cell components have been increasingly linked to the pathogenesis of the NDs and, in particular PD ^16,17^. The presence of the lipopolysaccharides (LPS), a major component of gram-negative bacteria and indication of the gut-brain axis dysfunction, in blood can activate microglia and eventually leads to chronic neuroinflammation. LPS can also promote α-synuclein aggregation, a hallmark of PD and its progression, which could also lead to further neurodegeneration and CI ^18^. Release of gram-positive bacterial components such as peptidoglycan and lipoteichoic acid, could stimulate immune responses and promote the secretion of proinflammatory cytokines and contribute to the neuroinflammation^19,20^.

Similarly, oral health of PD patients has also been shown to impact the course of disease ^21^. α-synuclein, the molecule that forms aggregates in neurons in PD, can be detected in saliva. The presence of α-synuclein in the oral cavity frequently results in reduced saliva production and difficulty swallowing ^22,23^, and report the association of NDs with dysphagia. Oral bacteria contribute to chronic inflammation and neurodegeneration through various mechanisms. Opportunistic pathogens in the oral cavity, which proliferate due to dysbiosis within the oral ecosystem, can form biofilms leading to bacterial overgrowth ^24–26^. These biofilms, often associated with gingivitis and periodontitis ^27^, enable bacteria to enter the bloodstream, potentially causing bacteraemia and systemic inflammation ^28^. *Porphyromonas gingivalis* is a well-studied oral pathogen and has been observed in Alzheimer’s disease (AD) brains and active periodontitis have been reported to impact CI ^29–31^, and in bacteraemia cases it can induce the blood-brain permeability barrier ^32^. The presence of inflammation, bacteraemia, and dysfunction of the mucosal barriers can lead to spontaneous dissemination of the bacteria across tissues ^33^. Simultaneously, the use of specific drugs, such as proton pump inhibitors and antibiotics, to treat stomach reflux, gastritis, and ulcers, that are common conditions in PD patients ^34^, modulate and accelerate the microbial translocation ^35^. The presence of oral pathobionts and their overgrowth, exacerbates gut dysbiosis and systematic inflammation, as has been reported in several other diseases ^36,37^.

In this study we used metagenomics of faeces and saliva in a cohort of PD patients with MCI (PD-MCI) or full dementia (PDD), together with a healthy control cohort. We hypothesize that compositional and functional differences in the microbiomes exist along the CI spectrum and that these differences, in turn, impact PD progression. We use a combination of machine learning approaches together with functional, correlative and network analyses associate microbiome changes with CI. Through this, we aim to establish an oral-gut-brain axis in PD to bring forth a more mechanistic understanding of the human microbiome in ND.

## RESULTS

### Gut and oral microbiome composition is associated with cognitive decline in Parkinson’s Disease

We performed shotgun metagenomics on 228 saliva and faecal samples taken from 41 Parkinson’s Disease (PD) patients with mild cognitive impairment (PD-MCI), 47 patients with full dementia (PDD) and 26 healthy controls (HC) (**Figure 1A, Supplementary Table S1**). The age and gender distribution between the PD-MCI and PDD groups were similar (**Table 1**, **Figure 1B**), with a mean age of 67.27 years (SD = 8.75 years) and 70.89 years (SD = 7.34) for PD-MCI and PDD, respectively, and 34.15% and 40.43% of females for PD-MCI and PDD, respectively. Cognitive assessment using the Mini-Mental State Examination (MMSE) and Clinical Dementia Rating Scale (CDRS) revealed a significant difference between PD-MCI and PDD patients (**Table 1**, **Figure 1C**). Additionally, motor function parameters, including the Unified Parkinson’s Disease Rating Scale (UPDRS), Hoehn and Yahr Scale (HYE), and disease duration, also showed significant differences between the two groups (**Table 1**, **Figure 1D**). Overall, these findings indicate that patients with MCI exhibit distinct cognitive and motor characteristics compared to those with full dementia.

**FIGURE 1.**
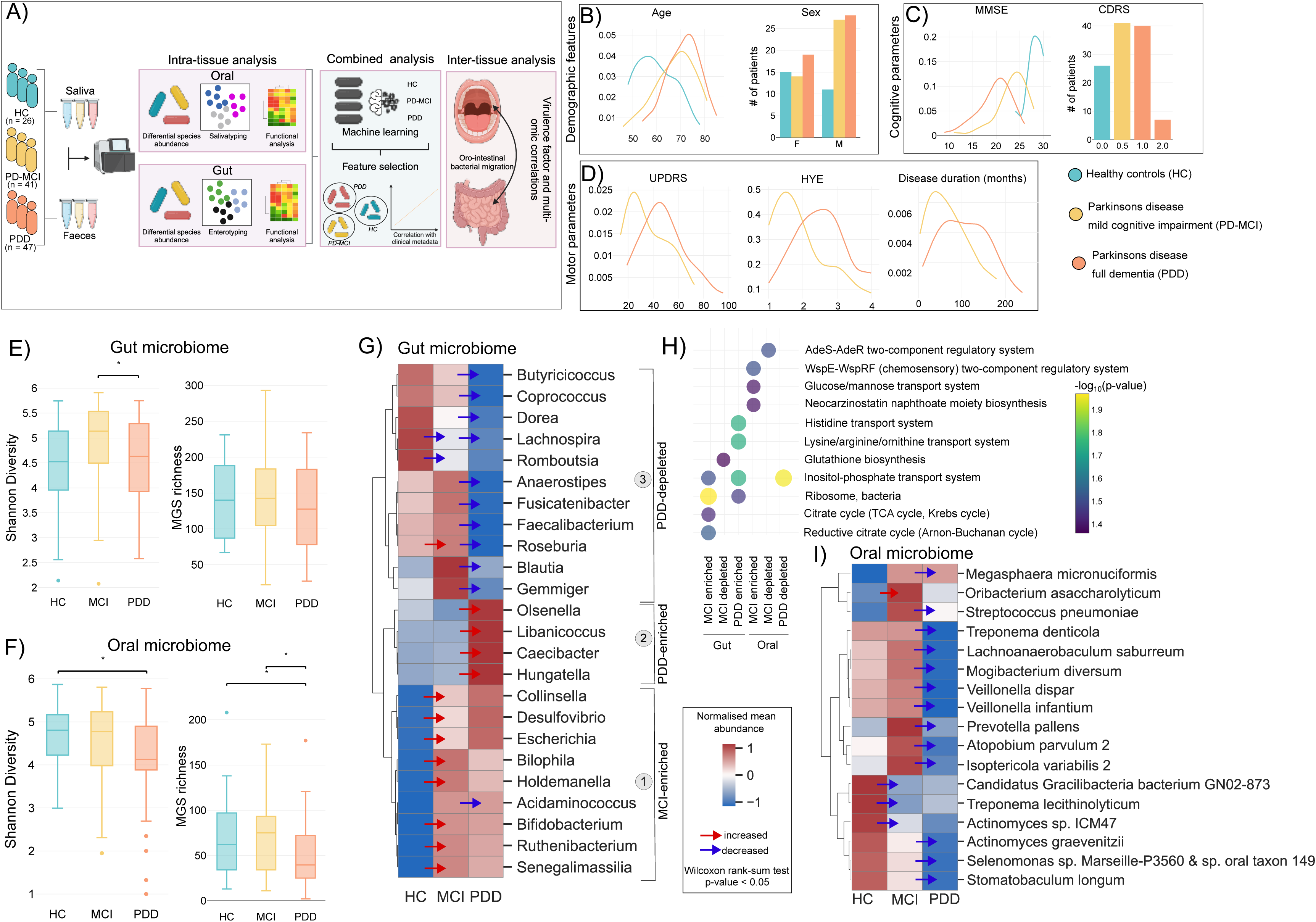
Gut and oral microbiome dysbiosis in Parkinson’s Disease patients with different degrees of cognitive impairment. A) **Study and methodology overview.** A total of 114 individuals were included in the study. This included 41 Parkinson’s disease (PD) patients with mild cognitive impairment (PD-MCI), 47 patients with full dementia (PDD) as well as 26 healthy controls (HC). Saliva and faecal samples were collected from all individuals and used for DNA extraction to perform shotgun metagenomics (**Methods**). We first performed intra-tissue analysis by investigating compositional and functional microbial changes in gut and oral separately. Thereafter, we combined gut and oral data to perform predictive modelling using machine learning. Finally, we investigated whether the translocation of oral species to the gut potentially impact disease. B) **Demographic features of study population.** Distribution of age and gender for HC, PD-MCI and PDD patients. C) **Key cognitive features of study population.** Distribution of scores for the Mini Mental State Examination (MMSE) and CDRS scores. D) **Key motor parameters of study population.** Distribution of UPDRS, HYE and disease duration (months). E) **Shannon diversity and MGS richness of the gut microbiome.** Significance was calculated with a Wilcoxon rank-sum test with an asterisk (*) indicating p-value < 0.05. F) **Shannon diversity and MGS richness of the oral microbiome.** Significance was calculated with a Wilcoxon rank-sum test with an asterisk (*) indicating p-value < 0.05. G) **Relative abundance changes of genera in the gut microbiome.** MGS were mapped to their corresponding genus and differentially abundant genera were calculated using Wilcoxon rank-sum test with a p-value cut-off of 0.05. Significantly changing genera were visualized using normalized mean abundance by calculating Z-scores for each genus. Arrows indicate either a significant depletion (blue) or increase (red) of the abundance of a genus. H) **Metabolic pathway enrichment of different patient populations.** The gene counts of all samples were used to map against the KEGG database to calculate genes counts for metabolic genes that were then used for enrichment analysis by first calculating differentially abundant genes using Wilcoxon rank-sum tests. Enrichment of KEGG modules were then performed using hypergeometric enrichment with a p-value cut-off of 0.05. I) **Relative abundance changes of genera in the oral microbiome.** MGS were mapped to their corresponding species and differentially abundant species were calculated using Wilcoxon rank-sum test with a p-value cut-off of 0.05. Significantly changing species were visualized using normalized mean abundance by calculating Z-scores for each species. Arrows indicate either a significant depletion (blue) or increase (red) of the abundance of a species.

**Table 1:** Demographic and clinical features of the study cohort.

| Table 1: Demographic and clinical features of the study cohort |  |  |  |
| --- | --- | --- | --- |
|  | HC | PD-MCI | PDD |
| *Demographic features* |  |  |  |
| Age (years, mean ± std) | 59.62 ± 8.30 | 67.27 ± 8.75 | 70.89 ± 7.34 |
| Gender (Female) | 15 (57.69%) | 14 (34.15%) | 19 (40.43%) |
| Education (years mean ± std) | 59.62 ± 8.30 | 67.27 ± 8.75 | 70.89 ± 7.34 |
| *Cognitive features* |  |  |  |
| MMSE (mean ± std) | 28.00 ± 1.85 | 23.34 ± 3.42 | 19.66 ± 3.58 |
| CDR (mean ± std) | 0.00 ± 0.00 | 0.50 ± 0.00 | 1.15 ± 0.36 |
| *Motor features* |  |  |  |
| UPDRS (mean ± std) | - | 35.59 ± 16.16 | 47.70 ± 17.60 |
| HYE (mean ± std) | - | 1.97 ± 0.90 | 2.54 ± 0.86 |
| PD duration (months, mean ± std) | - | 69.88 ± 49.67 | 105.20 ± 56.85 |

To study the in-depth compositional and functional changes of the gut and oral microbiomes on cognitive impairment and progression in Parkinson’s Disease (PD), we performed deep sequencing with an average of 31.9 reads and 28.2 million reads per sample, respectively, and quantified microbial genes and metagenomic species (MGSs) (**Figure 1A**). Stool and saliva samples were mapped to respective gene catalogues to satisfactory standard with and average mapping rate of 67.50% and 39.32%, respectively, which allowed us to confidently proceed with downstream analysis to identify MGSs (**Figure S1A, Supplementary Table S2**, **Supplementary Table S3, Supplementary Table S4**). Our analysis revealed a significant decrease in gut microbiome diversity in PDD patients compared to those with PD-MCI (**Figure 1E**) and a decrease in both diversity and MGS richness in the oral microbiome of PDD patients (**Figure 1F, Method**). Taxonomic profiling at the phylum level in gut and oral microbiome showed Actinobacteria was increased in the gut of PD-MCI and PDD patients while Bacteroidetes was decreased (**Figure S1B, Figure S1C, Supplementary Table S5**). In the oral cavity we found a decrease in Actinobacteria, Bacteroidetes, Firmicutes, Proteobacteria and Spirochaetes, specifically in PDD patients. These findings suggest that global alterations in both gut and oral microbiomes are present and may be linked to cognitive decline in PD.

To further investigate how the composition of the microbiome changes at varying levels of cognitive impairment, we performed differential abundance testing of MGSs together with functional enrichment analysis. In the gut microbiome, we identified three clusters of signatures characterized by distinctly different genera (**Figure 1G**). The first genera cluster showed the enrichment of species for *Bifidobacterium longum, Bilophila wadsworthia, Ruthenibacterium lactatiformans* in PD-MCI patients (**Figure S1D**). Notably, consistent with a previous report, *Desulfovibrio* genus also increases with PD severity^38^. These species were functionally enriched for energy generating metabolic pathways such as citrate cycle, as well inositol-phosphate metabolism but depleted for glutathione biosynthesis (**Figure 1H**). The second cluster represents a significant enrichment in PDD patients with opportunistic pathogen species from genera such as *Olsenella sp. Marseille-P2912* and *Hungatella* (**Figure S1D**) and with functional enrichment like those of PD-MCI but additionally enriched for several amino acid transport systems (**Figure 1H**). The third cluster represents a distinct depletion signature characteristic in PDD patients that predominantly consists of commensal and beneficial microbes. Several butyrate-producing microbes such as *Roseburia faecis, Faecalibacterium prausnitzii* together with several *Ruminococcus* species were all depleted in PDD compared to PD-MCI (**Figure 1G**, **Figure S1D**). In addition, compared to HC patients, PDD patients also show enrichment of *B. longum, B. adolescentis, R. lactatiformans* (**Figure S1E**). We then reconstructed a correlation network using the gut microbiome genera of these three clusters which showed that cluster 1 and cluster 2 have an overall negative correlation with cluster 3 further supporting that the depletion signature in PDD identified in cluster 3 (**Figure S1F**). As expected, genera in cluster 2 had a stronger negative correlation with cluster 3 compared to cluster 1 and cluster 3, however, we interestingly also observed a strong negative correlation between *Bifidobacterium* in cluster 1 and other genera in cluster 3 suggesting that *Bifidobacterium* is a strong driver in gut microbiome community structures and that its increase leads to depletion of other species.

In the oral cavity, we observed an overall depletion of several species in PD-MCI and PDD patients compared to HC, which could show the loss of diversity and commensalism in the oral cavity and opportunity for pathogens to triumph (**Figure S1G**). We did, however, find a significant increase in abundance of *Oribacterium asaccharolyticum* in PD-MCI (**Figure 1I**). Other potential pathogenic species such as *Streptococcus pneumoniae* and *Prevotella pallens* were also increased in PD-MCI compared to HC, albeit not significantly. Our functional enrichment showed that hexose sugar transport, chemosensory two component regulatory system and neocarzinostatin antibiotic biosynthesis are enriched in PD-MCI (**Figure 1H**).

### Enterotypes and salivatypes revealed distinct functional features for Parkinson’s disease and cognitive impairment

To understand the compositional changes of gut and oral communities we performed principal coordinate analysis (PCoA). Although there was a statistically significant separation in the gut (gut; PERMANOVA p-value=0.017, oral; PERMANOVA p-value=0.495), the clustering in both cases were discernible (**Figure S2A, Figure S2B**). Dirichlet multinomial mixture modelling has previously been shown to bring about hidden community structures in microbiome data that otherwise cannot clearly be distinguished with supervised clustering methods ^39^. Using this approach, we identified three clusters for gut microbiome, termed enterotypes, enriched for different bacterial genera (ENT1,2 and 3; **Figure 2A**). Our clinical study groups (PD-MCI and PDD) were enriched with different enterotypes (**Figure 2B**), and we found significant clustering of enterotypes using PCoA (**Figure 2C**). HC patients were enriched for ENT2 with a distinct signature of commensal *Prevotella* bacteria while PD-MCI and PDD showed a decrease in ENT2 instead. ENT2 was depleted for cytochrome C oxidase that potentially indicate deficient energy metabolism in the gut of PD patients (**Figure 2D**). PD-MCI patients were enriched for ENT1 that showed a signature for *Bacteroides* and *Alistipes*. PDD were enriched for *Bacteroides* of ENT1 and ENT3. ENT3 in PDD was depleted for aminoacyl-tRNA biosynthesis and ribosomal pathways suggests overall less translation of proteins and a reduction in citrate cycle pathways can potentially also indicate reduced production of SCFAs. In contrast, branch-chain amino acid (BCAA) production such as isoleucine was enriched which has been shown to be linked to different diseases including PD ^40–43^.

**FIGURE 2.**
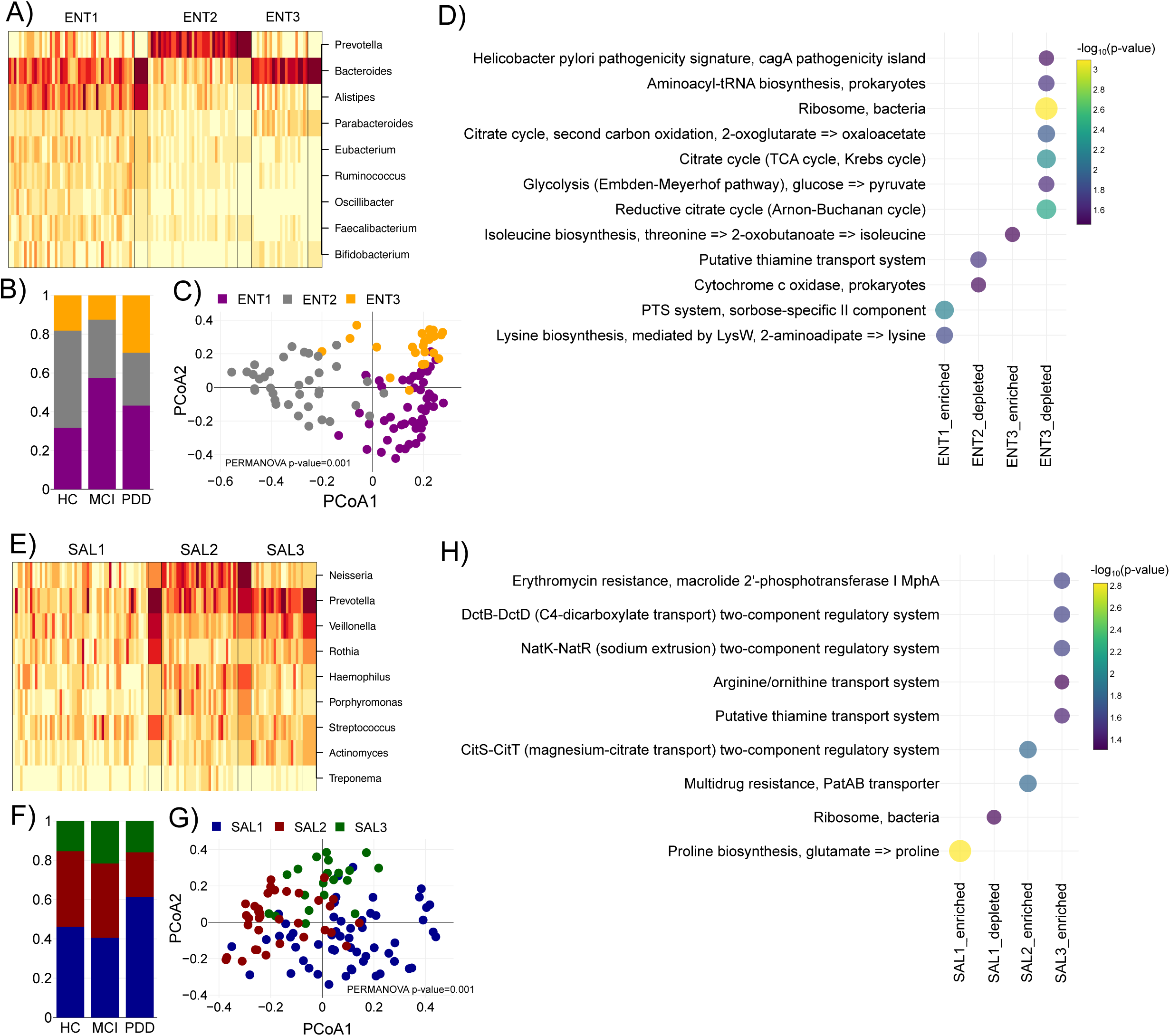
Enterotypes and salivatypes have distinct functional features. **A) Heatmap showing the genus abundance for three enterotypes (ET).** Using Dirichlet multinomial clustering we identified an optimal number of three clusters that differentiate the gut microbiome composition termed enterotype 1-3 (ENT1-3). Each cell in the heatmap depicts the relative abundance of a particular genus to each sample. **B)** Fraction of samples classified as ENT1-3 for HC, PD-MCI and PDD, respectively. **C) Principal coordinate analysis (PCoA) of gut samples.** The Bray-Curtis distance between all samples were calculated using species abundances and then used to perform PCoA. Each sample in the PCoA plot was coloured according to its assigned enterotype. **D)** Functional enrichment of KEGG modules for ENT1-3 **E) Heatmap showing the genus abundance for three salivatypes (SAL).** Using Dirichlet multinomial clustering we identified an optimal number of three clusters that differentiate the oral microbiome composition termed salivatype 1-3 (SAL1-3). Each cell in the heatmap depicts the relative abundance of a particular genus to each sample. **F)** Fraction of samples that were classified as SAL1-3 for HC, PD-MCI and PDD, respectively. **G) Principal coordinate analysis (PCoA) of oral samples.** The Bray-Curtis distance between all samples were calculated using species abundances and then used to perform PCoA. Each sample in the PCoA plot was coloured according to its assigned salivatype. **H)** Functional enrichment of KEGG modules for SAL1-3

We then identified three clusters, termed salivatypes, in the oral cavity (SAL1, 2 and 3; **Figure 2E**). This result pointed out the PD-MCI and PDD groups were enriched to different salivatypes (**Figure 2F**), and salivatypes clustered significantly using PCoA (**Figure 2G**). SAL1, enriched in PDD patients, was increased in pathobionts such as *Streptococcus, Rothia* and *Veillonella* and showed an enrichment for proline biosynthesis. PDD patients also showed a depletion in SAL2 that were dominated by *Neisseria*. Interestingly, SAL2 were enriched for multidrug resistance and its depletion in PDD potentially indicate a dysfunctional microbial community.

### Gut and oral biomarkers accurately predict clinical phenotypes

The observation that patients can be stratified by their gut and oral microbiomes could reflect the potential to use the microbiome for prediction of clinical outcomes and further be extended to identify novel prognostic biomarkers. We therefore used the abundances of gut and oral microbial species together with clinical metadata as features for predicting clinical outcomes (PD-MCI and PDD) using two machine learning (ML) algorithms and then used SHapley Additive exPlanations (SHAP) scoring to interpret model predictions and explain the contribution of features, or species, towards model predictions.

In four different predictions, we used gut (SIM1), oral (SIM2), both (SIM3; gut and oral) and both together with age, gender, and education (SIM4) of these patients, as features for ML prediction. Using AUCROC and accuracy we showed that SIM4 performed the best compared to other models with an average AUCROC of 69.42% and average accuracy score of 66.91% (**Supplementary Table S6, Methods**). It was particularly interesting to see that the inclusion of clinical metadata (age, gender, and education) improved the AUCROC score. We therefore focussed further analysis and feature selection on outcomes of SIM4 that included gut and oral species abundances together with clinical metadata.

We first showed accurate prediction of PD-MCI compared to HC (AUCROC: SVC=0.84, RF=0.89; Accuracy: SVC=0.88, RF=0.77; **Figure 3A, Methods**) as well as PDD compared to HC (AUCROC: SVC=0.82, RF=0.86; Accuracy: SVC=0.73, RF=0.78; **Figure 3B**). Of particular interest was to assess whether the microbiome can be used to distinguish different levels of cognitive decline by comparing PD-MCI and PDD patients. It was therefore supportive to also accurately predict PDD compared to PD-MCI (AUCROC: SVC=0.59, RF=0.57; Accuracy: SVC=0.45, RF=0.5; **Figure 3C**). Furthermore, in predicting PDD versus PD-MCI the inclusion of clinical metadata in SIM4 improved the AUCROC and accuracy of the model compared to SIM3 where only gut and oral metagenomics was used (AUCROC: SIM3=0.41, SIM4=0.58; Accuracy: SIM3=0.395, SIM4=0.475; **Supplementary Table S6**). This indicates that microbial species changes are sensitive enough to differentiate between PD with varying clinical features, and the inclusion of additional features like age and gender can significantly improve model predictions.

**FIGURE 3.**
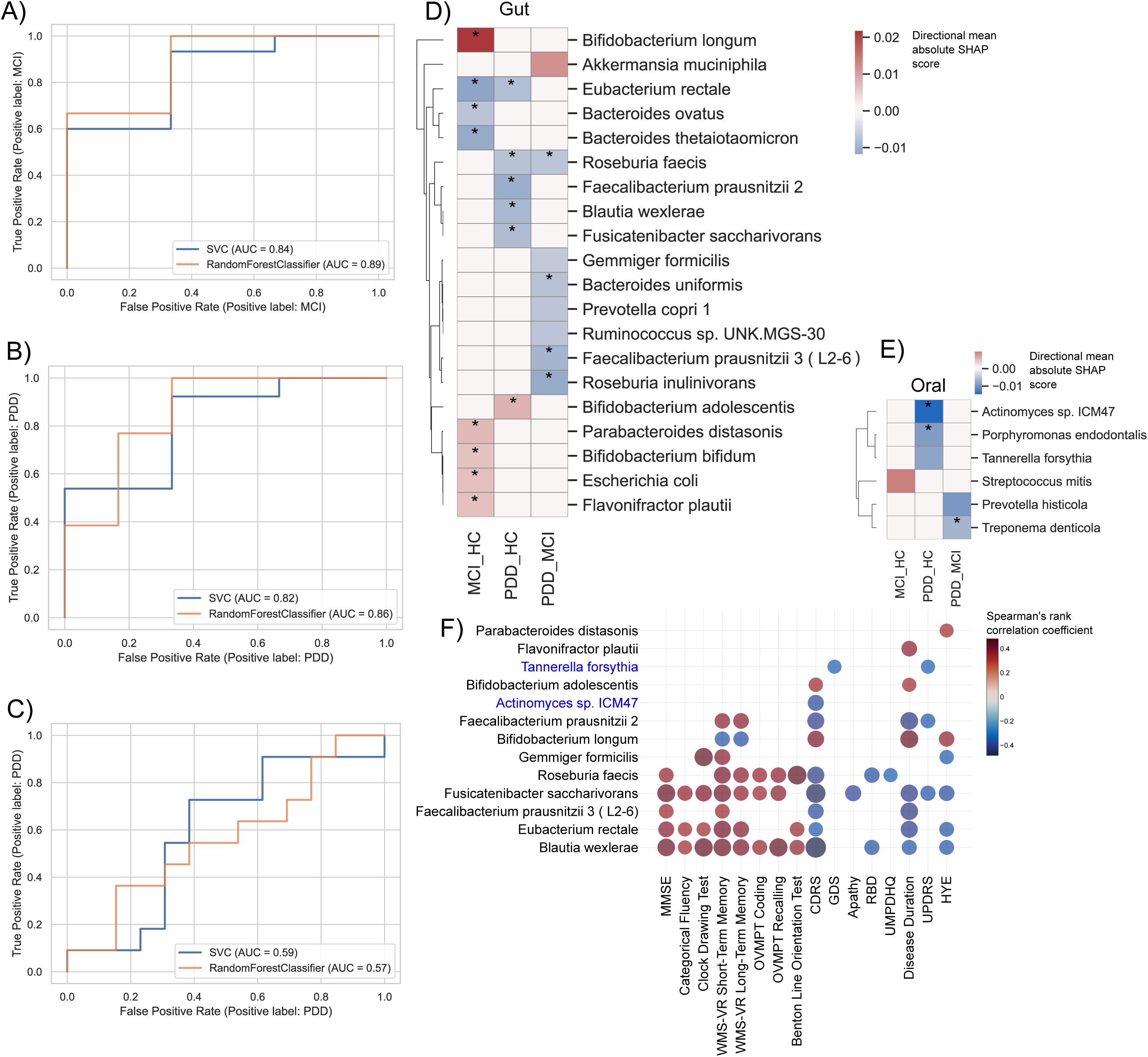
Machine learning models using gut and oral microbial species accurately predicts Parkinson’s Disease clinical phenotypes. **A-C) Random Forest (RF) and Support Vector Classification (SVC) models for prediction of clinical state.** Using the species abundances of gut and oral metagenomes as features, two machine learning models were built to predict PD-MCI versus HC (A), PDD versus HC (B) and PDD versus MCI (C). The ROC curves show good AUC for all models. **D-E) Top features contributing to RF prediction in three different models.** Using SHAP calculations we calculated the contribution of features (gut and oral microbial species, D and E, respectively) to the prediction of each model. The asterisks indicate whether these species were found to be significantly dysregulated using differential abundance analysis (Table S5). F) **Association of features to clinical metadata.** Model features (species abundances) was correlated with clinical metadata using spearman rank correlation. The significant associations (p-value < 0.01) are indicated with asterisks (*) and the colour is indicative of the correlation coefficient.

We then applied SHAP interpretation on the ML outputs to determine the contribution of species to model predictions (**Methods**). Several species that were significantly dysregulated (**Supplementary Table S5**) were also identified as important features for predicting PD-MCI and PDD. The decrease in beneficial bacteria such as *Faecalibacterium prausnitzii, Roseburia faecis, Roseburia inulinivorans, Eubacterium rectale* in the gut together with a decrease of *Treponema denticola, Porphyromonas endodontalis* and *Actinomyces* in the oral cavity were the most important features in predicting PDD (**Figure 3D**, **Figure 3E**). *Bacteroides uniformis* important for PDD prediction, have been shown to be associated with PD by increasing DAT (dopamine transporter) binding of dopamine and its decrease associated here with PDD might indicate poor dopamine metabolism that cause cognitive decline ^44^. It was also interesting to see that the increase in *Akkermansia muciniphila* that has previously been associated with PD ^2,12^ was an important feature for PDD prediction. The fact that the abundance of *A. muciniphila* did not significantly change in PDD using differential abundance analysis but with predictive modelling shows the value of using ML together with differential abundance. Another interesting observation was that species belonging to the *Bfidiobacterium* genus, also previously shown to be important in PD pathogenesis, were important features for predicting PD-MCI and PDD. However, *B. longum* and *B. bifidum* were associated with PD-MCI prediction while *B. adolescentis* was associated with PDD.

We finally associated top predictor species with clinical metadata and found several correlations (**Figure 3F**). For example, the abundance of *E. rectale*, *F. prausnitzii* and *R. faecis* showed a significant association with MMSE (**Figure S3A**). Other cognitive parameters also showed an overall positive correlation with these species. For example, *Fusicatenibacter saccharivorans* showed a positive correlation with categorical fluency tests of patients (**Figure S3B**). Motor parameters such as UPDRS, in addition, had correlation with oral pathogen *Tannerella forsythia* (**Figure S3C**).

### Oral microbiome in the gut is enriched for virulence factors that contributes to PD pathophysiology and cognitive decline

The translocation of oral-specific microbial species or features to the gut lumen, a phenomenon termed oralization of the gut, has previously been associated with different diseases ^45,46^. As mentioned in the introduction, the migration of bacteria, and even fragments of their genomes to other body sites and tissues ^47^, can increase the release of bacterial metabolites and cellular components causing systematic inflammation ^48^. To explore whether gut oralization is associated with cognitive impairment (CI), we mapped gut metagenomes against a non-redundant oral microbial gene catalogue to identify oral-specific genes in the gut (**Methods**).

After retrieving the gene counts from the cross-mapping gut samples, initially we performed gene richness analysis and observed PDD patients have significant enriched oral microbial genes in the gut (**Figure 4A**). To determine whether these genes potentially play a role in pathogenesis, we characterized potential virulence factors (VFs) and showed that PD-MCI and PDD patients have enriched oral-specific VFs in the gut compared to HC (**Figure 4B**). When calculating enriched or depleted VFs we found an overall enrichment of VFs in PD-MCI and PDD compared to HC and that 187 of enriched VFs overlap in PD-MCI and PDD (**Figure 4C, Supplementary Table S7**).

**FIGURE 4.**
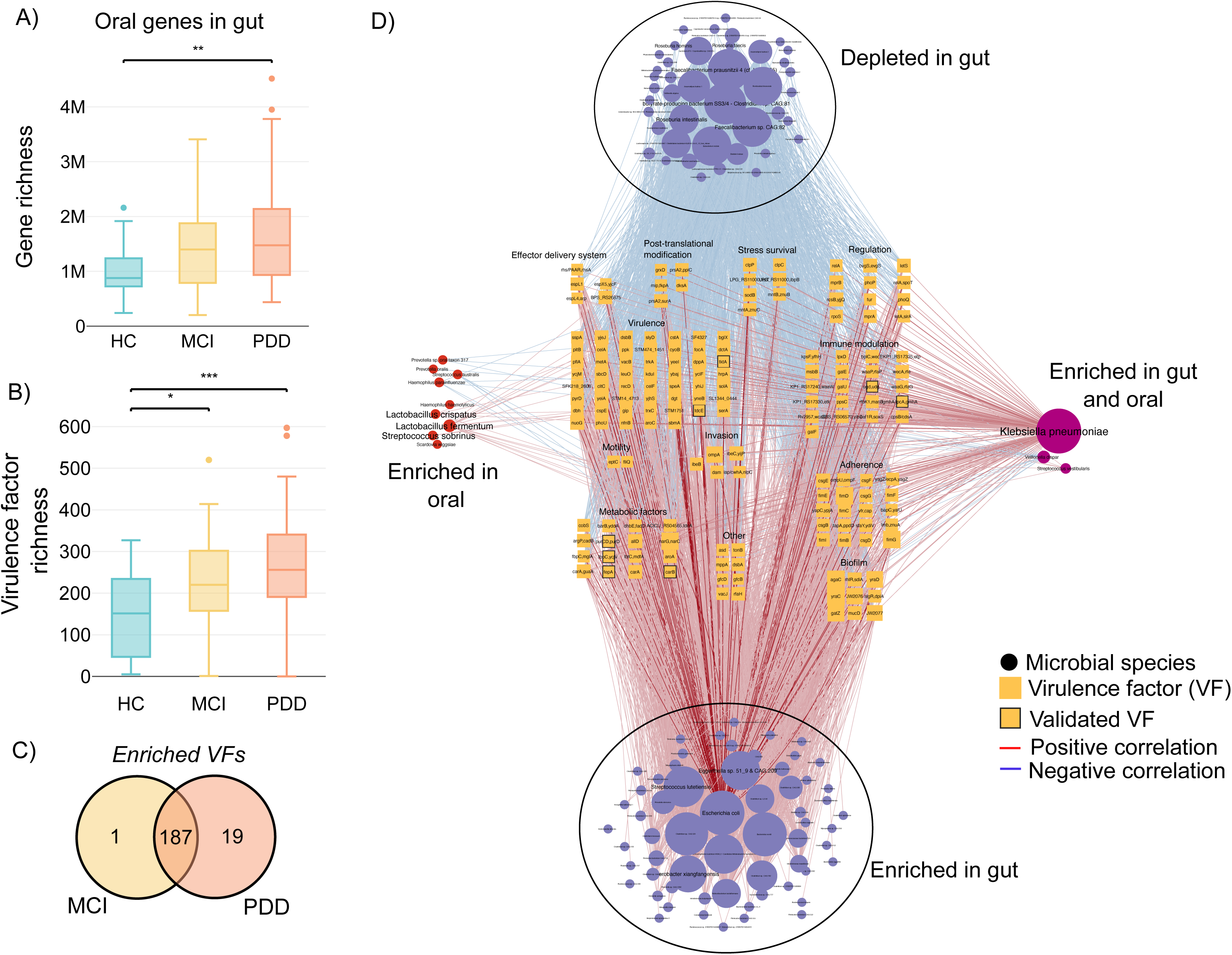
The sharing of gut and oral species is associated with increased virulence that contributes to disease progression. **A) Richness of oral-specific microbial genes in the gut.** All gut samples were mapped against the gene catalogue of oral microbial genes. The richness of genes was then calculated for different study groups. **B) Richness of oral-specific virulence factors (VFs) in the gut.** VFs were identified using sequence alignment of the oral genes against the PATRIC database (Methods). The richness of VFs was then calculated for different study groups. **C) Venn diagram of enriched VFs in PD-MCI and PDD patients**. Differentially abundant VFs were calculated for PD-MCI and PDD patients against HC, respectively, using Wilcoxon rank-sum tests (p-value 0.05). **D) Network analysis of enriched VFs in PD patients**. The 187 significantly enriched VFs were correlated with species in the gut and oral. Using network analysis, we then identified several enriched and depleted species in the gut that are positively and negatively correlated with these VFs, respectively. Virulence factors were aligned using BLASTP against saliva metaproteomics from the same subjects and significantly aligned VFs are annotated with black borders. Node sizes reflect the number of edges connected to the node.

To further understand how specific species contribute to enrichment of VFs we first identified which MGSs in both the gut and the oral contain these VFs within their pan-genomes (**Figure S4**). In both the gut and the oral, *Escherichia coli* contained the most VFs. In the oral cavity, other pathobionts such as *Klebsiella pneumonia*, *Cronobacter sakazakii* and *Streptococcus salivarius* also showed several VFs as part of their genomes. Similarly, in the gut, several *Klebsiella* and *Enterobacter* species were found enriched for VFs.

To elucidate underlying community structures, we constructed an integrative correlation network between gut species, oral species, and oral VFs in the gut (**Figure 4D**). Using network analysis, we identified two clusters of co-occurring species in the gut. The first consisted of enriched species that all had a positive correlation with VFs while the second consisted of depleted species that all had a negative correlation with VFs. This was an interesting observation because enriched species were primarily pathobionts such as *E. coli*, *Egerthella sp.* and *Enterobacter xiangfangensis* which were most connected, while depleted species were all species associated with a healthier gut such as *Feacalibacerium spps* and *Roseburia intestinalis*. Overall, this suggested that the enrichment of pathogenic species in PD, which are associated with increased VFs, potentially cause a depletion of commensal bacteria, reduces species diversity, and gut barrier dysfunction. We also found that *K. pneumonia* was positively correlated in both the gut and oral with VFs and that *Streptococcus sobrinus, Lactobacillus fermentum* and *L. crispatus* were enriched in the oral and positively correlated with VFs. The significant positive correlations between the oral and gut enriched species with virulence in the PDD and PD-MCI.

Enriched VFs had different functions related to stress, immune modulation, adherence, biofilm formation, invasion, and metabolism (**Figure 4D, Supplementary Table S8**). The most connected VFs, *agaC*, *yraC*, *yraD, fimG, fimD*, *narG*, *narC*, *gatZ* and *ompA*, were all involved in biofilm formation, adherence, and invasion. Outer membrane protein A (*ompA*) contributes to brain microvascular endothelial cells (BMECs) invasion via a ligand-receptor interaction. Further investigation revealed that *ibeB*, *ibeC and lmb* are also involved in brain microvascular endothelial cell invasion. It was also interesting to find that several VFs related to immune modulation are involved in LPS synthesis.

We then used saliva proteomics previously generated for the same cohort of patients ^26^ to identify whether any of the discovered oral VFs in gut metagenomes were enriched in the oral cavity that could potentially suggest that these VFs have translocated to the gut and serve as a validation (**Figure 4D**). We were able to validate eight VFs using this approach. D-sedoheptulose 7-phosphate isomerase (*gmhA*), is a major immunogen involved in LPS biosynthesis and UDP-glucose 6-dehydrogenase (*udg*) also involved in immune modulation assist in the evasion of the host immune system by protecting bacteria from opsonophagocytosis and serum killing (**Supplementary Table S8**). Other validated VFs were involved in metabolism and two genes, outer membrane receptor for ferric enterobactin and colicins B, D (*fepA*) and ATP-binding protein (*fbpC*), were responsible for iron uptake by bacterial cells. Lastly, carbamoyl-phosphate synthase large chain (*carB*) mediates bacterial resistance to reactive oxygen species (ROS) and is important for phagosomal escape.

Together our integrative analysis results showed a potential novel mechanism for how gut microbiome dysbiosis contribute to PD pathogenesis and specifically CI (**Figure 5**). The enrichment of pathobionts and depletion of butyrate-producing commensals in the gut lumen are accompanied by the infiltration of VFs from the oral microbiome that, establishing an oral-gut axis that disrupt community structure. The formation of biofilms in the gut lumen can bring forth two potential mechanistic actions of how the oral-gut dysbiosis establish an axis with the brain and contribute to disease (**Figure 5**). In the first instance, VFs such as *ompA, ibeB / C* and *lmb* can directly interact with brain endothelial cells if secreted to the bloodstream. In the second instance, VFs annotations revealed several functions related to survival and replication and importantly increased lipopolysaccharide production (LPS). LPS can also contribute to a direct action through increase of gut-wall permeability. Overall, indirect actions give rise to dysfunctional host immunity that can impact disease progression.

**FIGURE 5.**
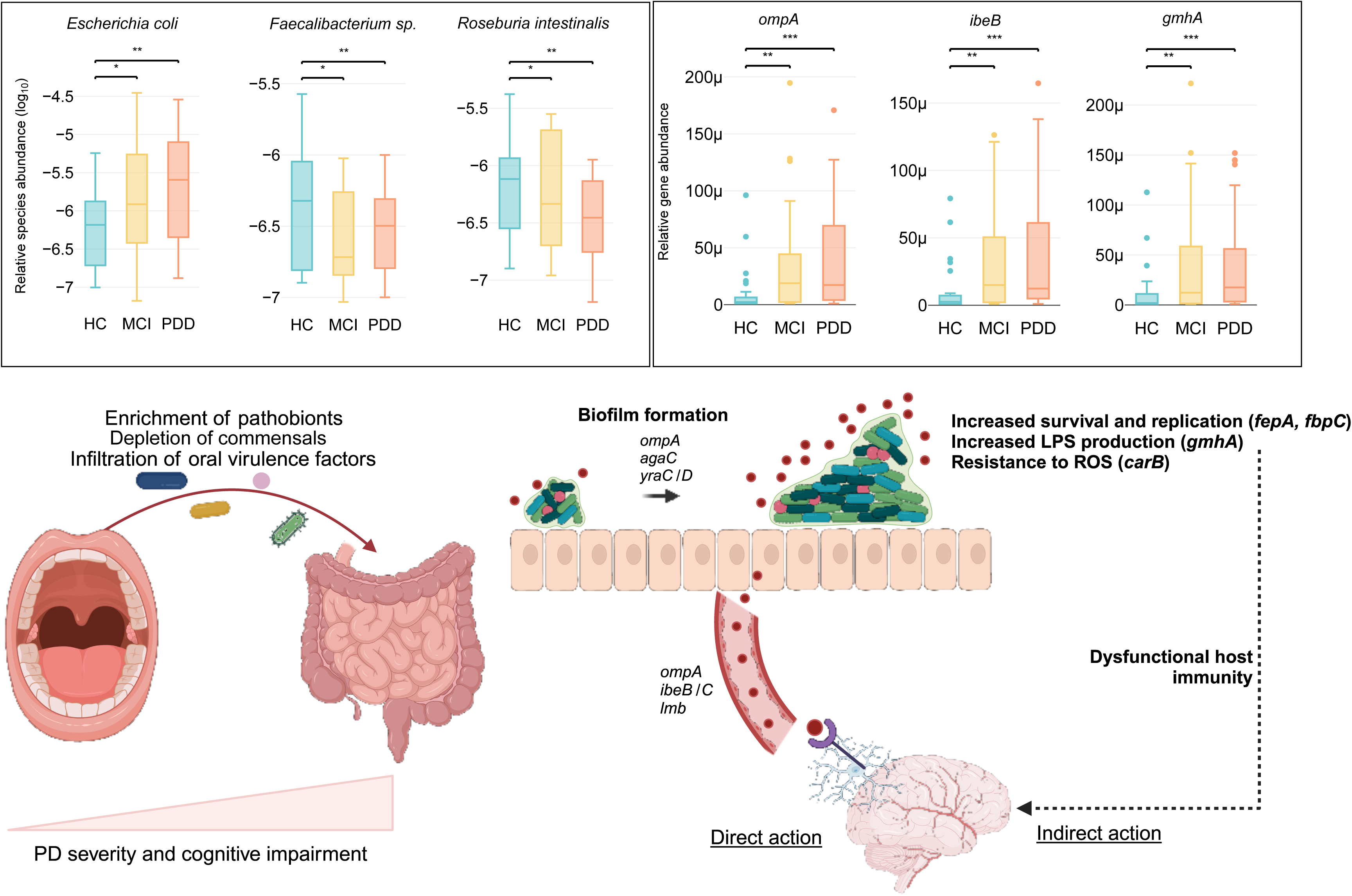
An oral-gut-brain axis established through the infiltration of oral virulence factors to the gut. Network analysis revealed key pathobiont species, such as *E. coli,* enriched in the gut while commensal species such as *Faecalibacterium* and *R. intestinalis* are decreasing. Together with the increase of VFs such as *ompA, ibeB* and *gmhA* we propose a new oral-gut-brain axis where these compositional changes promote biofilm formation in the gut that, in turn, increase production of VFs that directly interact with the brain. We also propose an indirect mechanism where increased bacterial survival and replication, LPS production and bacterial protection from ROS, could lead to dysfunctional host immunity.

## DISCUSSION

The human gut and oral microbiomes have each been implicated to play a role in the pathology of PD and other neurodegenerative diseases ^2^. Our study reveals a significant connection between the oral and gut microbiomes in PD patients, emphasizing the functional role of this oral-gut microbiome continuum in cognitive impairment. Most notably, our functional analysis shows that infiltration of oral microbiome in the gut are responsible for increased virulence factors in the gut lumen of PD patients.

### Microbes previously associated with PD change with CI

The increase of microbial species of the *Bifidobacterium, Lactobacillus* and *Akkermansia* genera in the gut have consistently been shown as features of PD ^2,11,12,25,45^. At the same time, butyrate producers, *Roseburia, Faecalibacterium* and *Blautia* are consistently decreased in PD patients. Our results show, for the first time, that CI of PD patients impacts these features if we consider different stages of CI.

We found that *B. longum* is enriched in PD-MCI and PDD while *B. adolescentis* are only significantly enriched in PDD, suggesting that it is important to consider species specific changes with relation to CI. *Bifidobacterium adolescentis* can therefore be a potential marker for cognitive decline in PD. Similarly, *L. mucosae* was enriched in PD-MCI and PDD while *L. salivarius* and *L. gasseri* were only enriched in PDD and *L. johnsonii* was only enriched in MCI. Although not as frequently reported, *Ruthenibacterium lactatiformans*, enriched in PD-MCI and PDD, were also previously shown associated with PD ^11^.

In congruence with previous studies, we found several *Faecalibacterium* spp, *Roseburia* spp, and *Blautia* spp depleted in PDD ^2,11^. We also identified other butyrate producers such as *Ruminococcus sp.* to be depleted. Interestingly these species were depleted in PDD but not in PD-MCI compared to HC. Furthermore, we found some of these species, e.g., *Faecalibacterium prausnitzii* and *Blautia wexlerae*, were also significantly depleted in PDD compared to PD-MCI. Taken together, these results therefore suggest that depletion of species that are generally associated with a healthy gut environment decline as a function of CI and that they could potentially be used as markers of CI in PD.

There are fewer studies and less consensus on the compositional changes in the oral microbiome in PD. However, the recent discovery of *Porphyromonas gingivalis*, an oral pathogen causing periodontitis, in the brains of patients with Alzheimer’s disease patients brought forth an interest on the functional role of the oral microbiome and CI ^29^. We found that *P. endodontalis* is significantly enriched in PDD. *Porphyromonas endodontalis* is also found in patients with chronic periodontitis and its abundance is correlated with *P. gingivalis* ^49^. We therefore show for the first time another species of *Porphyromonas* that are associated with CI and hypothesize that is could potentially play a functional role in PD pathogenesis.

### A combination of unsupervised and supervised ML methodologies reveals functional features and associations with clinical data

Several studies have explored applying different ML methods using microbiome data ^50^. We have used both an unsupervised and supervised approach and subsequently used model predictions for functional insights and feature selection.

Unsupervised clustering has previously been used in metagenomics to establish a defined compositional signature of microbiome, termed enterotypes (ENT) ^39^. The concept of the three distinct enterotypes initially proposed has been challenged ^51^ and therefore we performed a *de novo* enterotype assignment for each sample. The depletion of a *Prevotella* enterotype and an enrichment of a *Bacteroides* enterotype in the gut of PD patients recapitulated our previous study ^12^. Both PD-MCI and PDD showed a *Bacteroides* enterotype, however, PD-MCI also showed an *Alistipes* enterotype, a genus showed to be highly elevated in PD ^11^ and specifically enriched in PD-MCI ^6^. Therefore, our findings suggest that *Alistipes* could be a marker for differentiation of CI in PD. An interesting observation was the enrichment of isoleucine biosynthesis is PDD. Branch-chain amino acid (BCAA) metabolism in the gut has been linked to several diseases including correlation between BCAA levels and PD clinical symptoms ^40,52^. Using the approach of unsupervised clustering to identify salivatypes (SAL) has not been performed before in PD. Most notably, the enrichment of SAL1 in PDD because of increased *Prevotella*, *Veilonella, Rothia* and *Streptococcus* have enrichment of proline biosynthesis. Dysregulated proline metabolism has been shown in neuronal dysfunction and psychiatric disorders and in particular a recent study has functionally linked proline metabolism and gut microbiome in depression ^53,54^.

Several studies have explored using microbiome data as input for ML classifiers and more complex ML models ^45,50^. A major challenge, however, is development of robust interpreter methods to identify the most important features that contribute to model prediction. Here, we have used SHAP that is a state-of-the-art method for this purpose that we have used in a previous study ^45^. We showed that combining gut and oral microbiome data significantly improves model predictions, which is important for establishing clinically feasible biomarkers for PD. We found that the increase of *A. muciniphila*, commonly increased in PD, was an important feature for prediction of PDD against PD-MCI, however, *A. muciniphila* was not significantly increased in PDD when performing differential abundance analysis. This highlights the value that interpretable ML models can add to existing microbiome methodologies by identifying trends across the dataset that contribute to a specific disease phenotype where the species does not necessarily significantly change in abundance between groups. Given the significant influence of diet on microbiome, systematic investigation of dietary nutrient ^55–57^ on microbiome composition and functions could elucidate a more mechanistic role of *A. muciniphila* in this context.

The depletion of *Bacteroides uniformis* was an important feature for differentiating between PDD and PD-MCI. The dopamine transporter (DAT) is responsible for transport of dopamine, the most common treatment for patients with PD. In a recent study, Hartstra *et al* showed that faecal microbiota transplant of *B. uniformis* increased DAT binding and importantly that the gut-brain axis can be modulated ^58^. Our results here therefore shows that this species might be particularly important for cognition through dopamine metabolism and that its depletion leads to increased cognitive decline.

### The oralisation of the gut lumen correlates with increased virulence

There have been several studies that have shown that the translocation of oral bacterial species, often opportunistic pathogens, to the gut lumen drive disease phenotypes ^45,59,60^. Although this provides novel avenues for biomarkers, the underlying mechanisms and functionality in particular diseases remain largely unclear.

The enrichment of oral VFs in the gut of PD patients, in particular PDD, shows that these species potentially exert specific functions in the gut. The in-depth network analysis also highlighted underlying microbiome community structures where enriched pathogenic species such as *Escherichia coli, Enterobacter xiangfangensis* and *Egerthella* supress commensal butyrate producing species such as *Faecalibacterium spp, Roseburia intestinalis* and *Roseburia faecis*. Competition between commensals and pathogenic bacteria and an imbalance are known to contribute to disease and here were show that it correlates with increased production of VFs by pathobionts providing mechanistic insights that could be exploited therapeutically.

Our integrative allowed us to propose a mechanistic oral-gut-brain axis mediated by increased production of VFs in the gut (**Figure 5**). In the first instance, several oral VFs are involved in biofilm formation and adherence. The formation of biofilms on the outer mucosal layer can lead to mucosal invasion by bringing bacteria close to the epithelium that contribute to a leaky gut. *OmpA*^61^, highly connected in our network, is a key VFs that mediate the formation of bacterial biofilms but has interestingly also been shown to be a contributor to invasion of brain microvascular endothelial cells (BMECs) *via* ligand-receptor interaction. Together, with the increased connectivity of genes with similar function, for example *ibeB* that has been shown to invade BMECs ^62^, our findings suggest a mechanism where infiltration of oral factors to the gut could cross the gut wall and interact with brain endothelial cells.

We hypothesize that the formation of biofilms in the gut contributes to overall virulence and dysregulation of the immune system through various mechanisms. Firstly, we validated two oral VFs, *fepA* and *fbpC*, involved in iron acquisition and iron metabolism that has been shown to be important in the pathogenicity and survival of pathogens such as *E. coli* ^63^. In addition, reactive oxygen species (ROS) that are produced as by-products of metabolism can induce DNA damage and mediation through increased *carB* potentially protects biofilm formation. Finally, the production of lipopolysaccharide (LPS) has been shown to contribute to several diseases and a leaky gut. Apart from *gmhA* that was validated here, several other oral VFs in the gut were related to LPS metabolism. Together, we therefore hypothesize that the infiltration of the oral microbiome to the gut creates a dysregulated microbial community structure, that promotes a leaky gut, increase pathogen survival which in turn increases LPS production and other VFs that can interact with BMECs either directly or indirectly to promote CI and PD.

Our study provides compelling evidence that the interplay between oral and gut microbiomes significantly influences Parkinson’s Disease pathology and cognitive impairment. The translocation of oral microbial species to the gut, along with their associated virulence factors, highlights new avenues for understanding disease mechanisms and developing potential biomarkers and therapeutic avenues. The integration of machine learning techniques and microbiome data enhances our ability to identify key functional features and underscores the importance of a multifaceted approach in advancing our knowledge of PD and related neurodegenerative diseases.

## METHODS

### Study subjects and clinical characteristics

The study was approved by the ethics committee of the Istanbul Medipol University with authorization number 10840098-604.01.01-E.3958, and informed consent was obtained from all participants. A total of 114 subjects (HC = 26; PD-MCI = 41; PDD = 47) were recruited at two tertiary training hospitals including the Medipol Training and Research Hospital in the neurology clinic and Bakirkoy Research and Training Hospital for Psychiatric and Neurological Diseases. Participants were part of a larger cohort recruited into an ongoing prospective study on CI in PD. Clinical and demographic information, including age, gender, years of education were collected at clinic visits (**Supplementary Table S1**). The patients were examined by experienced neurologists and the diagnosis of PD was made within the framework of the “United Kingdom Parkinson’s Disease Society Brain Bank” criteria. Subjects with previous head trauma, stroke, or exposure to toxic substances, substance abuse, history of antibiotic or probiotic use within last one-month, chronic severe diseases (diabetes, cancer, kidney failure, etc.), autoimmune diseases, smokers, and those with symptoms suggestive of Parkinson’s plus syndromes were excluded from the study. The Hoehn-Yahr Stages Parkinson’s Staging Scale was utilized to assess the disease stage, while The Movement Disorder Society’s diagnostic criteria for Parkinson’s Disease Dementia were employed for evaluating dementia ^64^. The diagnosis of Mild Cognitive Impairment (MCI) was established following the guidelines outlined by Litvan et al. ^65^, employing level II criteria. This involved conducting a thorough cognitive assessment utilizing the MDS task force diagnostic criteria, which comprises neuropsychological evaluations covering two tests for each of the five cognitive domains.

### Sample preparation and metagenomics sequencing

A total of 228 samples (114 stool and 114 saliva) (HC = 26, PD-MCI = 41, PDD = 47) were used for shotgun metagenomics sequencing. Microbial DNA was extracted from saliva samples using the DNeasy PowerSoil kit (Qiagen, Hilden, Germany) with previously described modifications ^26^. For stool samples, the same extraction kit was used with adjustments to the manufacturer’s protocol. Stool samples were transferred to the PowerBead tube and homogenized by bead-beating using a Next Advance Bullet Blender (30 s at level 4, 30 s incubation on ice, and 30 s at level 4). Subsequently, the manufacturer’s protocol was followed without further modification. Shotgun sequencing libraries were prepared according to Illumina’s Nextera XT library preparation protocol and sequenced using a NovaSeq600 platform with a 2 × 150 paired-end kit. All stool samples passed quality control and a total of 107 saliva samples passed quality control (HC = 26, PD-MCI = 37, PDD = 44).

### Microbial gene and metagenomics species (MGS) quantification

Raw sequencing reads were mapped and counted using the METEOR pipeline (available at: https://github.com/sysbiomelab/meteor_pipeline). For gut samples, the IGC2 gene catalogue ^66^ of human gut microbiome was used as reference and for oral samples the HS_8.4_oral gene catalogue ^67^ was used. Mapping was performed using a >95% identity threshold to account for gene variability and the non-redundant nature of the catalogue (**Supplementary Table S2**). This generated gene count matrices that were then subjected to downsizing and normalization (reads per kilo base per million mapped reads (RPKM method) to generate the gene frequency matrix for downstream analysis. For the gut samples, downsizing was done at 5 million reads before normalization to correct for differences in sequencing depth. After downsizing a total of 106 samples were remaining (HC = 22, PD-MCI = 40, PDD = 44) and for the oral samples, only normalization was performed (**Figure S1A, Supplementary Table S2**). The resulting gene matrices were then projected on previously reconstructed metagenomic species (MGSs) using the top 50 marker genes per MGS to calculate MGS abundances for each sample (**Supplementary Table S3**). Analysis was performed using the R package MetaOMineR (momr) designed to analyse large quantitative metagenomics datasets ^68^.

### Functional gene annotation and analysis

The normalised gene count matrices were annotated for functional investigation. Amino acid sequences of the gut and oral catalogues were aligned to amino acid sequences of KEGG orthologs (KEGG database version 82) ^69^ using Diamond (version 0.9.22.123) ^70^ and best hit alignments with e-value ≤ 10^−5^ and bit scores ≥ 60 were considered. Amino acid sequences of the gut and oral catalogues were aligned to amino acid sequences of proteins in the PATRIC database ^71^ using BLASTP and best hit alignments with e-value ≤ 10^−10^ and identity of > 80% were considered. The virulence factor database (VFDB) incorporates the PATRIC database and gives more in depth-annotations and descriptions ^72^. We therefore enhanced the PATRIC databases annotations with that of VFDB. This was then used to calculate gene abundances for metabolic genes and virulence factors. For enriched and depleted KEGG modules we first identified differentially abundant metabolic genes using Wilcoxon rank-sum tests. For comparison of clinical study groups in Figure 1H we used a p-value cut-off of 0.05 and for comparison between enterotypes and salivatypes (**Figure 2D and Figure 2H**) we used a p-value cut-off of 0.01. For both analyses only genes with a log_2_foldchange of 2 or more were considered. We then used these genes to identify significantly enriched or depleted KEGG modules using hypergeometric tests adjusted for false discovery using the Benjamini-Hochberg procedure and considered p-adjusted < 0.05 as significantly changing modules.

### Richness, diversity analysis and differential abundance analysis

Richness was calculated as the sum of the number of MGS per sample and Shannon diversity was calculated using the *skbio* package in Python. Beta-diversity was done by first calculating the Bray-curtis distance between all samples using the *distance* function in scipy and then performing principal coordinate analysis using the *skbio* package. For differential abundance analysis the calculated abundances of MGSs were mapped to different taxonomical ranks (phylum, class, order, family, genus, or ^73^species) and the sum of each taxon calculated per sample. We then used Wilcoxon rank-sum tests with false discovery rate adjustment using the Benjamini-Hochberg procedure to calculate differentially abundant microbes at these different taxonomical ranks. Details of statistical cut-offs can be found in the text.

### Machine learning classification to predict clinical outcomes

We used the Scikit-learn python package to train random forest (RF) and support vector classification (SVC) models to predict different clinical outcomes ^74^. Training and testing were performed on randomly selected samples split 70% and 30% of the full dataset, respectively, with a fixed random seed to ensure the reproducibility of the model. The following hyperparameters were set for the RF model: ‘random_state’: 1, ‘n_estimators’: 500, ‘bootstrap’: True and for the SVC model: ‘random_state’: 1. All other parameters were kept as their default. Model performance was measured using AUROC scoring and accuracy. Python implementation of the explainable AI algorithm, Shapley Additive ExPlanations (SHAP), was used to show the feature (species) contribution to disease classification the mean absolute SHAP score for each disease predictive model was determined using the sign of the Spearman rank correlation between the feature value and the SHAP score. Positive values indicate that a higher relative abundance is more likely to classify the disease than in healthy samples. Negative values indicate that a lower relative abundance is more likely to classify the disease than in healthy samples.

### Integrative correlation network analysis of VFs and MGSs

We first mapped the gut metagenomes against the oral gene catalogue to identify oral specific genes in the gut. Enriched and depleted VFs were calculated using Wilcoxon rank-sum tests (p-value < 0.05) considering a positive and negative log-fold change, respectively. We then performed a Spearman’s rank correlation between enriched VFs and MGSs in the gut and oral, respectively, using a p-value cut-off of 0.01. The protein sequences of enriched VFs were aligned to previously generated saliva metaproteomics using BLASTP with an e-value cut-off of 10^−7^ and percentage identity of higher than 50. The integrative network was then visualised in Cytoscape and used to calculate the degree of connections of each node.

## Data availability

The metagenome data sequenced for this study from the total of 221 samples (114 stool and 107 saliva) can be found in the European Nucleotide Archive under the study accession PRJEB79944.

## Acknowledgments

This study was supported by grants from Engineering and Physical Sciences Research Council (EPSRC), EP/S001301/1, Biotechnology Biological Sciences Research Council (BBSRC) BB/S016899/1 and Science for Life Laboratory (SciLifeLab) to Saeed Shoaie and from the Scientific and Technological Research Council of Turkey (TÜBITAK) (grant no. 315S301) to Suleyman Yildirim. We acknowledge the Swedish National Infrastructure for Computing at SNIC through Uppsala Multidisciplinary Center for Advanced Computational Science (UPPMAX) under Project SNIC 2020-5-222, SNIC 2019/3-226, SNIC 2020/6-153 and King’s College London computational infrastructure facility, CREATE, for high performance computing.

## Author contributions

Conception and design, S.S., S.Y., and L.H., clinical metadata and sample collection and processing, T.K.D., Z.Y., M.A., N.H.Y., A.S., and L.H.; data analysis, F.C.; data interpretation, F.C., S.Y., S.S. A.M., M.U; manuscript writing—original draft, F.C.; review and editing, F.C., S.Y., S.S, M.A., A.M., M.U. All authors read and approved the final manuscript.

## REFERENCES

1 Global Burden of Disease Study, C. Global, regional, and national incidence, prevalence, and years lived with disability for 301 acute and chronic diseases and injuries in 188 countries, 1990-2013: a systematic analysis for the Global Burden of Disease Study 2013. Lancet 386, 743–800 (2015). 10.1016/S0140-6736(15)60692-4

2 Tan, A. H., Lim, S. Y. & Lang, A. E. The microbiome-gut-brain axis in Parkinson disease - from basic research to the clinic. Nat Rev Neurol 18, 476–495 (2022). 10.1038/s41582-022-00681-2

3 Rosario, D. et al. Systems Biology Approaches to Understand the Host-Microbiome Interactions in Neurodegenerative Diseases. Front Neurosci 14, 716 (2020). 10.3389/fnins.2020.00716

4 Kalia, L. V. & Lang, A. E. Parkinson’s disease. Lancet 386, 896–912 (2015). 10.1016/S0140-6736(14)61393-3

5 Aarsland, D. et al. Parkinson disease-associated cognitive impairment. Nat Rev Dis Primers 7, 47 (2021). 10.1038/s41572-021-00280-3

6 Ren, T. et al. Gut Microbiota Altered in Mild Cognitive Impairment Compared With Normal Cognition in Sporadic Parkinson’s Disease. Front Neurol 11, 137 (2020). 10.3389/fneur.2020.00137

7 O’Callaghan, C. & Lewis, S. J. G. Cognition in Parkinson’s Disease. Int Rev Neurobiol 133, 557–583 (2017). 10.1016/bs.irn.2017.05.002

8 Shi, H. et al. A fiber-deprived diet causes cognitive impairment and hippocampal microglia-mediated synaptic loss through the gut microbiota and metabolites. Microbiome 9, 223 (2021). 10.1186/s40168-021-01172-0

9 Olson, C. A. et al. Alterations in the gut microbiota contribute to cognitive impairment induced by the ketogenic diet and hypoxia. Cell Host Microbe 29, 1378–1392 e1376 (2021). 10.1016/j.chom.2021.07.004

10 Hill-Burns, E. M. et al. Parkinson’s disease and Parkinson’s disease medications have distinct signatures of the gut microbiome. Mov Disord 32, 739–749 (2017). 10.1002/mds.26942

11 Wallen, Z. D. et al. Metagenomics of Parkinson’s disease implicates the gut microbiome in multiple disease mechanisms. Nat Commun 13, 6958 (2022). 10.1038/s41467-022-34667-x

12 Rosario, D. et al. Systematic analysis of gut microbiome reveals the role of bacterial folate and homocysteine metabolism in Parkinson’s disease. Cell Rep 34, 108807 (2021). 10.1016/j.celrep.2021.108807

13 Shandilya, S., Kumar, S., Kumar Jha, N., Kumar Kesari, K. & Ruokolainen, J. Interplay of gut microbiota and oxidative stress: Perspective on neurodegeneration and neuroprotection. J Adv Res 38, 223–244 (2022). 10.1016/j.jare.2021.09.005

14 Loh, J. S. et al. Microbiota-gut-brain axis and its therapeutic applications in neurodegenerative diseases. Signal Transduct Target Ther 9, 37 (2024). 10.1038/s41392-024-01743-1

15 Alpino, G. C. A., Pereira-Sol, G. A., Dias, M. M. E., Aguiar, A. S. & Peluzio, M. Beneficial effects of butyrate on brain functions: A view of epigenetic. Crit Rev Food Sci Nutr 64, 3961–3970 (2024). 10.1080/10408398.2022.2137776

16 Tufekci, K. U., Genc, S. & Genc, K. The endotoxin-induced neuroinflammation model of Parkinson’s disease. Parkinsons Dis 2011, 487450 (2011). 10.4061/2011/487450

17 Zakaria, R. et al. Lipopolysaccharide-induced memory impairment in rats: a model of Alzheimer’s disease. Physiol Res 66, 553–565 (2017). 10.33549/physiolres.933480

18 Couch, Y., Alvarez-Erviti, L., Sibson, N. R., Wood, M. J. & Anthony, D. C. The acute inflammatory response to intranigral alpha-synuclein differs significantly from intranigral lipopolysaccharide and is exacerbated by peripheral inflammation. J Neuroinflammation 8, 166 (2011). 10.1186/1742-2094-8-166

19 Mayerhofer, R. et al. Diverse action of lipoteichoic acid and lipopolysaccharide on neuroinflammation, blood-brain barrier disruption, and anxiety in mice. Brain Behav Immun 60, 174–187 (2017). 10.1016/j.bbi.2016.10.011

20 Ginsburg, I. Role of lipoteichoic acid in infection and inflammation. Lancet Infect Dis 2, 171–179 (2002). 10.1016/s1473-3099(02)00226-8

21 Figura, M. & Friedman, A. In search of Parkinson’s disease biomarkers - is the answer in our mouths? A systematic review of the literature on salivary biomarkers of Parkinson’s disease. Neurol Neurochir Pol 54, 14–20 (2020). 10.5603/PJNNS.a2020.0011

22 Adler, C. H. & Beach, T. G. Neuropathological basis of nonmotor manifestations of Parkinson’s disease. Mov Disord 31, 1114–1119 (2016). 10.1002/mds.26605

23 Mu, L. et al. Alpha-Synuclein Pathology in Sensory Nerve Terminals of the Upper Aerodigestive Tract of Parkinson’s Disease Patients. Dysphagia 30, 404–417 (2015). 10.1007/s00455-015-9612-7

24 Mihaila, D. et al. The oral microbiome of early stage Parkinson’s disease and its relationship with functional measures of motor and non-motor function. PLoS One 14, e0218252 (2019). 10.1371/journal.pone.0218252

25 Jo, S. et al. Oral and gut dysbiosis leads to functional alterations in Parkinson’s disease. NPJ Parkinsons Dis 8, 87 (2022). 10.1038/s41531-022-00351-6

26 Arikan, M. et al. Metaproteogenomic analysis of saliva samples from Parkinson’s disease patients with cognitive impairment. NPJ Biofilms Microbiomes 9, 86 (2023). 10.1038/s41522-023-00452-x

27 Santamaria, P. et al. Microbiological and molecular profile of furcation defects in a population with untreated periodontitis. J Clin Periodontol (2024). 10.1111/jcpe.14034

28 Hajishengallis, G. & Chavakis, T. Local and systemic mechanisms linking periodontal disease and inflammatory comorbidities. Nat Rev Immunol 21, 426–440 (2021). 10.1038/s41577-020-00488-6

29 Dominy, S. S. et al. Porphyromonas gingivalis in Alzheimer’s disease brains: Evidence for disease causation and treatment with small-molecule inhibitors. Sci Adv 5, eaau3333 (2019). 10.1126/sciadv.aau3333

30 Sparks Stein, P., et al. Serum antibodies to periodontal pathogens are a risk factor for Alzheimer’s disease. Alzheimers Dement 8, 196–203 (2012). 10.1016/j.jalz.2011.04.006

31 Iwasaki, M. et al. Periodontitis, periodontal inflammation, and mild cognitive impairment: A 5-year cohort study. J Periodontal Res 54, 233–240 (2019). 10.1111/jre.12623

32 Lei, S. et al. Porphyromonas gingivalis bacteremia increases the permeability of the blood-brain barrier via the Mfsd2a/Caveolin-1 mediated transcytosis pathway. Int J Oral Sci 15, 3 (2023). 10.1038/s41368-022-00215-y

33 Fine, R. L., Manfredo Vieira, S., Gilmore, M. S. & Kriegel, M. A. Mechanisms and consequences of gut commensal translocation in chronic diseases. Gut Microbes 11, 217–230 (2020). 10.1080/19490976.2019.1629236

34 Makaroff, L., Gunn, A., Gervasoni, C. & Richy, F. Gastrointestinal disorders in Parkinson’s disease: prevalence and health outcomes in a US claims database. J Parkinsons Dis 1, 65–74 (2011). 10.3233/JPD-2011-001

35 Horvath, A. et al. Biomarkers for oralization during long-term proton pump inhibitor therapy predict survival in cirrhosis. Sci Rep 9, 12000 (2019). 10.1038/s41598-019-48352-5

36 Patel, V. C. et al. Rifaximin-alpha reduces gut-derived inflammation and mucin degradation in cirrhosis and encephalopathy: RIFSYS randomised controlled trial. J Hepatol 76, 332–342 (2022). 10.1016/j.jhep.2021.09.010

37 Nagao, J. I. et al. Pathobiont-responsive Th17 cells in gut-mouth axis provoke inflammatory oral disease and are modulated by intestinal microbiome. Cell Rep 40, 111314 (2022). 10.1016/j.celrep.2022.111314

38 Murros, K. E., Huynh, V. A., Takala, T. M. & Saris, P. E. J. Desulfovibrio Bacteria Are Associated With Parkinson’s Disease. Front Cell Infect Microbiol 11, 652617 (2021). 10.3389/fcimb.2021.652617

39 Arumugam, M. et al. Enterotypes of the human gut microbiome. Nature 473, 174–180 (2011). 10.1038/nature09944

40 Zhang, Y. et al. Plasma branched-chain and aromatic amino acids correlate with the gut microbiota and severity of Parkinson’s disease. NPJ Parkinsons Dis 8, 48 (2022). 10.1038/s41531-022-00312-z

41 Shoaie, S. et al. Quantifying Diet-Induced Metabolic Changes of the Human Gut Microbiome. Cell Metab 22, 320–331 (2015). 10.1016/j.cmet.2015.07.001

42 Proffitt, C. et al. Genome-scale metabolic modelling of the human gut microbiome reveals changes in the glyoxylate and dicarboxylate metabolism in metabolic disorders. iScience 25, 104513 (2022). 10.1016/j.isci.2022.104513

43 Ezzamouri, B. et al. Metabolic modelling of the human gut microbiome in type 2 diabetes patients in response to metformin treatment. NPJ Syst Biol Appl 9, 2 (2023). 10.1038/s41540-022-00261-6

44 Hamamah, S., Aghazarian, A., Nazaryan, A., Hajnal, A. & Covasa, M. Role of Microbiota-Gut-Brain Axis in Regulating Dopaminergic Signaling. Biomedicines 10 (2022). 10.3390/biomedicines10020436

45 Lee, S. et al. Global compositional and functional states of the human gut microbiome in health and disease. Genome Res 34, 967–978 (2024). 10.1101/gr.278637.123

46 Kitamoto, S. et al. The Intermucosal Connection between the Mouth and Gut in Commensal Pathobiont-Driven Colitis. Cell 182, 447–462 e414 (2020). 10.1016/j.cell.2020.05.048

47 Kataria, R., Shoaie, S., Grigoriadis, A. & Wan, J. C. M. Leveraging circulating microbial DNA for early cancer detection. Trends Cancer 9, 879–882 (2023). 10.1016/j.trecan.2023.08.001

48 Omenetti, S. et al. The Intestine Harbors Functionally Distinct Homeostatic Tissue-Resident and Inflammatory Th17 Cells. Immunity 51, 77–89 e76 (2019). 10.1016/j.immuni.2019.05.004

49 Lombardo Bedran, T. B., et al. Porphyromonas endodontalis in chronic periodontitis: a clinical and microbiological cross-sectional study. J Oral Microbiol 4 (2012). 10.3402/jom.v4i0.10123

50 Asnicar, F., Thomas, A. M., Passerini, A., Waldron, L. & Segata, N. Machine learning for microbiologists. Nat Rev Microbiol 22, 191–205 (2024). 10.1038/s41579-023-00984-1

51 Knights, D. et al. Rethinking “enterotypes”. Cell Host Microbe 16, 433–437 (2014). 10.1016/j.chom.2014.09.013

52 Wang, W. et al. Interactions between gut microbiota and Parkinson’s disease: The role of microbiota-derived amino acid metabolism. Front Aging Neurosci 14, 976316 (2022). 10.3389/fnagi.2022.976316

53 Yao, Y. & Han, W. Proline Metabolism in Neurological and Psychiatric Disorders. Mol Cells 45, 781–788 (2022). 10.14348/molcells.2022.0115

54 Mayneris-Perxachs, J. et al. Microbiota alterations in proline metabolism impact depression. Cell Metab 34, 681–701 e610 (2022). 10.1016/j.cmet.2022.04.001

55 Clasen, F. et al. Systematic diet composition swap in a mouse genome-scale metabolic model reveals determinants of obesogenic diet metabolism in liver cancer. iScience 26, 106040 (2023). 10.1016/j.isci.2023.106040

56 Bidkhori, G. et al. The Reactobiome Unravels a New Paradigm in Human Gut Microbiome Metabolism. bioRxiv, 2021.2002.2001.428114 (2021). 10.1101/2021.02.01.428114

57 Bidkhori, G. & Shoaie, S. MIGRENE: The toolbox for microbial and individualized GEMs, reactobiome and community network modelling. bioRxiv, 2023.2009.2001.555866 (2023). 10.1101/2023.09.01.555866

58 Hartstra, A. V. et al. Infusion of donor feces affects the gut-brain axis in humans with metabolic syndrome. Mol Metab 42, 101076 (2020). 10.1016/j.molmet.2020.101076

59 Lee, S. et al. Pathogenic entero-and salivatypes harbour changes in microbiome virulence and antimicrobial resistance genes with increasing chronic liver disease severity. bioRxiv, 2023.2008. 2006.552152 (2023).

60 Lee, S. et al. Transient colonizing microbes promote gut dysbiosis and functional impairment. NPJ Biofilms Microbiomes 10, 80 (2024). 10.1038/s41522-024-00561-1

61 Nie, D. et al. Outer membrane protein A (OmpA) as a potential therapeutic target for Acinetobacter baumannii infection. J Biomed Sci 27, 26 (2020). 10.1186/s12929-020-0617-7

62 Wang, Y. & Kim, K. S. Role of OmpA and IbeB in Escherichia coli K1 invasion of brain microvascular endothelial cells in vitro and in vivo. Pediatr Res 51, 559–563 (2002). 10.1203/00006450-200205000-00003

63 Ratledge, C. & Dover, L. G. Iron metabolism in pathogenic bacteria. Annu Rev Microbiol 54, 881–941 (2000). 10.1146/annurev.micro.54.1.881

64 Emre, M. et al. Clinical diagnostic criteria for dementia associated with Parkinson’s disease. Mov Disord 22, 1689–1707; quiz 1837 (2007). 10.1002/mds.21507

65 Litvan, I. et al. Diagnostic criteria for mild cognitive impairment in Parkinson’s disease: Movement Disorder Society Task Force guidelines. Mov Disord 27, 349–356 (2012). 10.1002/mds.24893

66 Li, J. et al. An integrated catalog of reference genes in the human gut microbiome. Nat Biotechnol 32, 834–841 (2014). 10.1038/nbt.2942

67 Le Chatelier, E., et al. A catalog of genes and species of the human oral microbiota. Portail Data INRAE 10 (2021).

68 Le Chatelier, E. et al. Richness of human gut microbiome correlates with metabolic markers. Nature 500, 541–546 (2013). 10.1038/nature12506

69 Kanehisa, M., Goto, S., Kawashima, S., Okuno, Y. & Hattori, M. The KEGG resource for deciphering the genome. Nucleic Acids Res 32, D277–280 (2004). 10.1093/nar/gkh063

70 Buchfink, B., Xie, C. & Huson, D. H. Fast and sensitive protein alignment using DIAMOND. Nat Methods 12, 59–60 (2015). 10.1038/nmeth.3176

71 Gillespie, J. J. et al. PATRIC: the comprehensive bacterial bioinformatics resource with a focus on human pathogenic species. Infect Immun 79, 4286–4298 (2011). 10.1128/IAI.00207-11

72 Liu, B., Zheng, D., Zhou, S., Chen, L. & Yang, J. VFDB 2022: a general classification scheme for bacterial virulence factors. Nucleic Acids Res 50, D912–D917 (2022). 10.1093/nar/gkab1107

73 Lundberg, S. M. & Lee, S.-I. A unified approach to interpreting model predictions. Advances in neural information processing systems 30 (2017).

74 Abraham, A. et al. Machine learning for neuroimaging with scikit-learn. Front Neuroinform 8, 14 (2014). 10.3389/fninf.2014.00014

