## Supplementary figures and images for "Microbiome signatures of virulence in the oral-gut-brain axis influence Parkinson’s disease and cognitive decline pathophysiology"

### Supplemental Figures

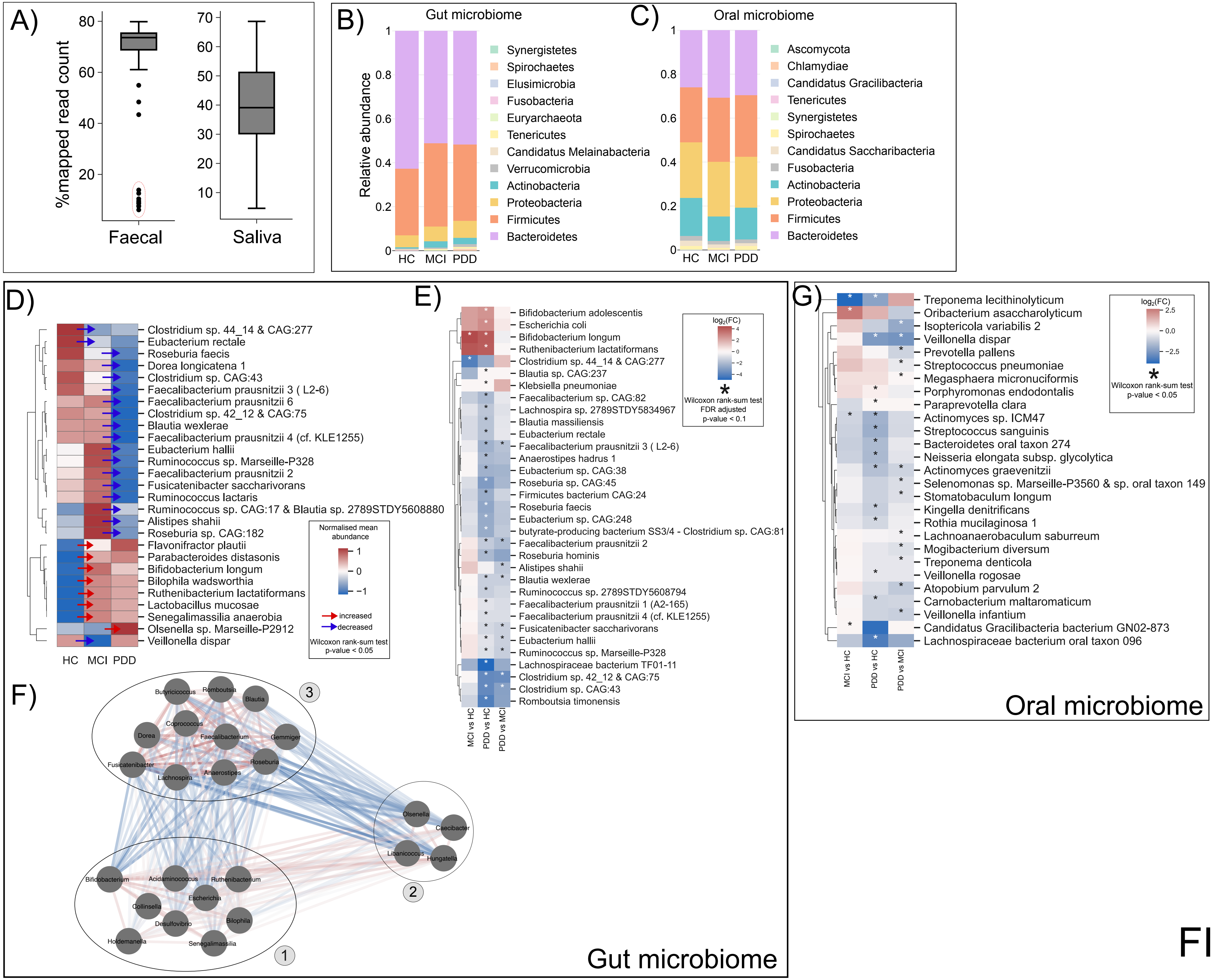

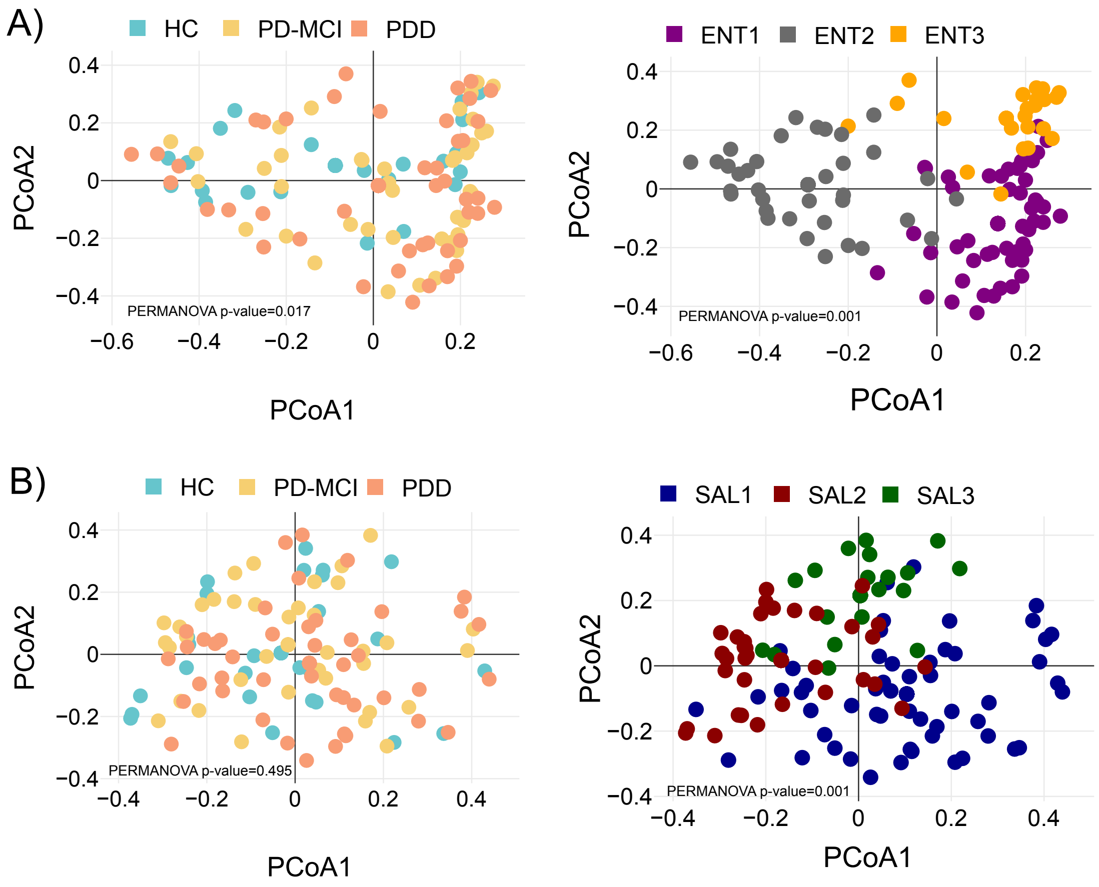

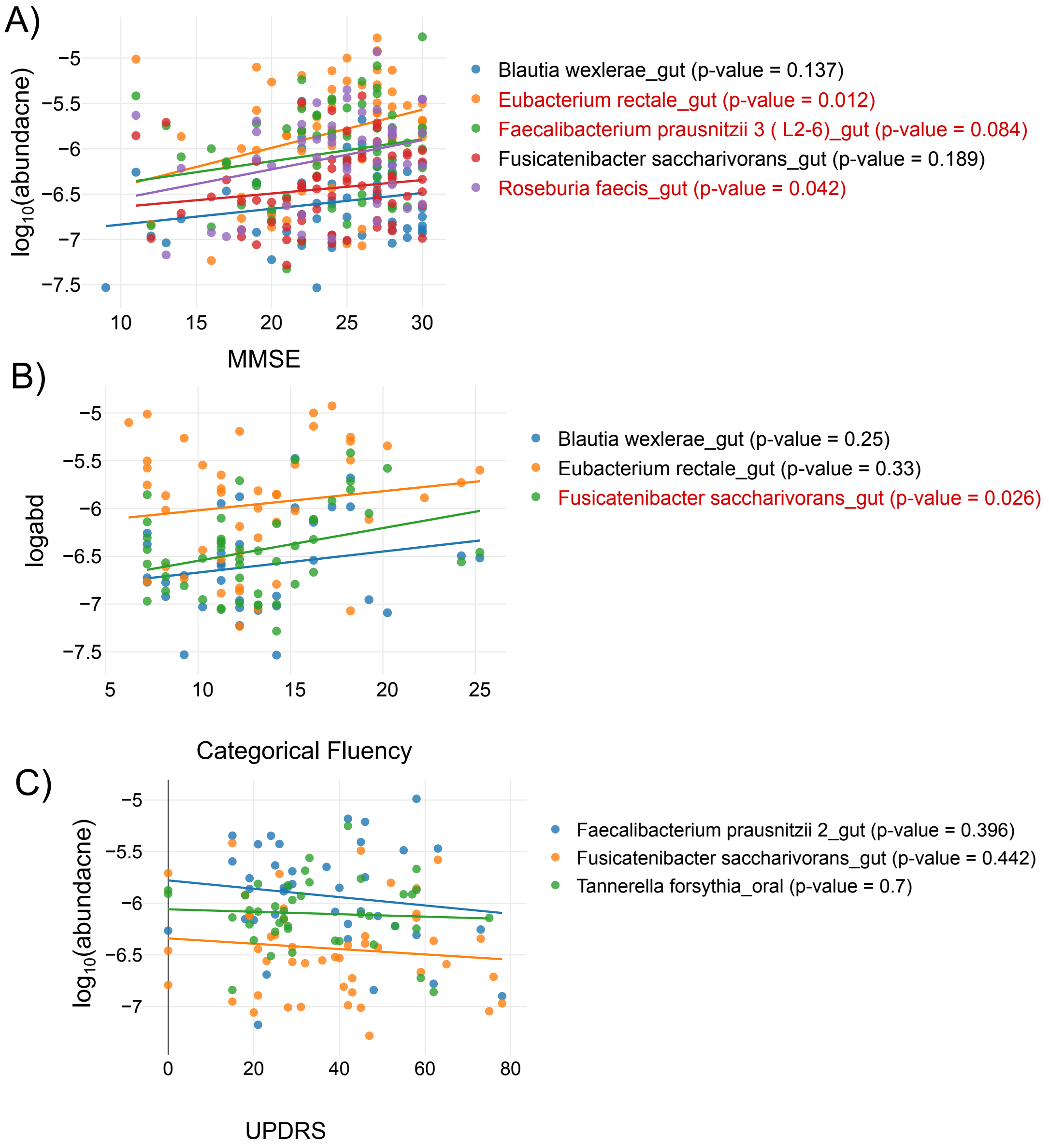

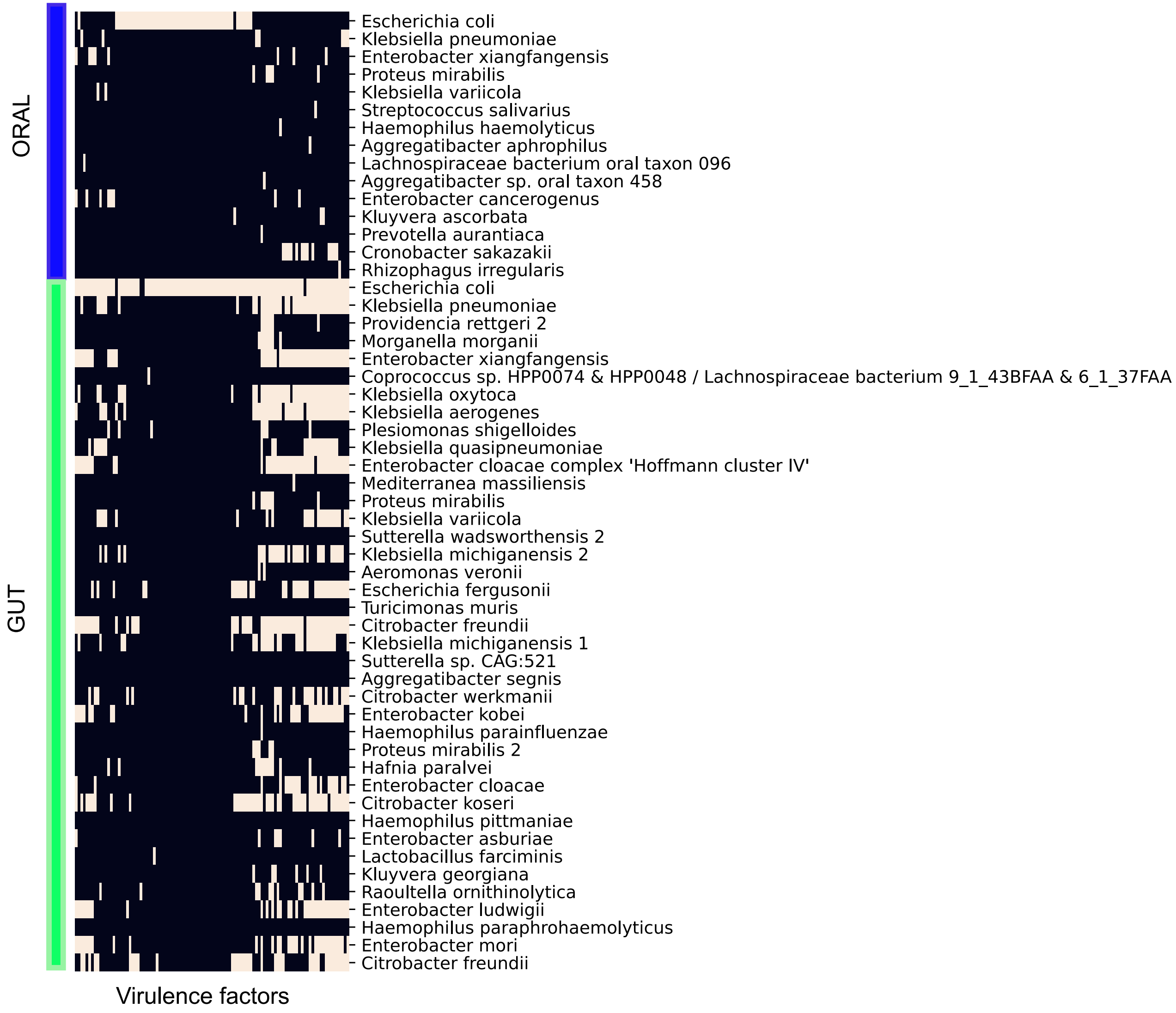
